# Weighted burden analysis of exome-sequenced late onset Alzheimer’s cases and controls provides further evidence for a role for *PSEN1* and suggests involvement of the PI3K/Akt/GSK-3β and WNT signalling pathways

**DOI:** 10.1101/596007

**Authors:** David Curtis, Kaushiki Bakaya, Leona Sharma, Sreejan Bandyopadhyay

## Abstract

Previous studies have implicated common and rare genetic variants as risk factors for late onset Alzheimer’s disease (AD, LOAD). Here, weighted burden analysis was applied to over 10,000 exome sequenced subjects from the Alzheimer’s Disease Sequencing Project. Analyses were carried out to investigate whether rare variants predicted to have a functional effect within a gene were more commonly seen in cases or in controls. Confirmatory results were obtained for *TREM2, ABCA7* and *SORL1*. Additional support was provided for *PSEN1* (p = 0.0002), which previously had been only weakly implicated in LOAD. There was suggestive evidence that functional variants in *PIK3R1, WNT7A, C1R* and *EXOC5* might increase risk and that variants in *TIAF1* and/or *NDRG2* might have a protective effect. Overall, there was strong evidence (p = 5 × 10^−6^) that variants in tyrosine phosphatase genes reduce the risk of developing LOAD. Since *PIK3R1* variants are expected to impair PI3K/Akt/GSK-3β signalling while variants in tyrosine phosphatase genes would enhance it, these findings are in line with those from animal models suggesting that this pathway is protective against AD.

## Introduction

A recent genetic meta-analysis of late onset Alzheimer’s Disease (AD, LOAD) to study the effects of common variants implicated Aβ, tau, immunity and lipid processing and reported a total of 25 genome-wise significant loci, including variants at or near TREM2, ABCA7 and SORL1 (Kunkle et al., 2019). Analyses of risk genes and pathways showed a general enrichment for rare variants but it was not possible to identify which specific variants were driving this enrichment. The Alzheimer’s Disease Sequencing Project (ADSP) has the aim of identifying rare genetic variants which increase or decrease the risk of Late Onset Alzheimer’s Disease (LOAD) and a full description of its membership, support and methodologies is provided on its website (https://www.niagads.org/adsp/content/about) (Beecham et al., 2017). A number of analyses utilising this dataset have been published to date. One study incorporating this dataset with others identified 19 loss of function variants of *SORL1* in cases as opposed to 1 in controls and a collapsing test of loss of function ultra-rare variants highlighted other genes including *GRID2IP, WDR76* and *GRN* (Raghavan et al., 2018). A subsequent study used variant-based and gene-based tests of association and followed up significant or suggestive results in other samples to implicate three novel genes, *IGHG3, AC099552.4* and *ZNF655* as well as novel and predicted functional genetic variants in genes previously associated with Alzheimer’s disease (AD) (Bis et al., 2018). In the ADSP samples alone, five genes were exome-wide significant: *ABCA7, TREM2, CBLC, OPRL1* and *GAS2L2*. Additionally, three individual variants were exome-wide significant: rs75932628 in *TREM2* (p = 4.8 × 10^−12^), rs2405442 in *PILRA* (p = 1.7 × 10^−7^), and a novel variant at 7:154,988,675 *AC099552.4* (p = 1.2 × 10^−7^). Another study focussed on a panel of 35 known dementia genes using variants thought likely to be consequential based on ClinVar, literature review and predicted effect (Blue et al., 2018). Using gene-based analyses in the ADSP case-control sample, *TREM2* was significant at p = 1.8 × 10^−11^ and *APOE* was significant at p = 0.0069 while *ARSA, CSF1R, PSEN1* and *MAPT* were less strongly implicated although were still significant at p < 0.05. In another study, 95 dementia associated genes were examined and it was reported that overall there was an excess of deleterious rare coding variants (Patel et al., 2019). Additionally, 10 cases and no controls carried rs149307620, a missense variant in *NOTCH3*, and 4 cases and no controls carried rs104894002, a high-impact variant in *TREM2* which in homozygous form causes Nasu-Hakola disease, which is a rare presenile dementia associated with recurrent bone fractures.

A limitation of these previous studies is that they focus on specific genes and/or types of variant. We have previously described a method for a carrying out a weighted burden analysis of all variants within a gene, assigning more weights to variants which are rare and to variants predicted to have a functional effect (Curtis, 2012; Curtis and UK10K Consortium, 2016). This method was applied to the ADSP sample in the hope that it would more completely capture the effect of variants influencing susceptibility to LOAD.

## Methods

Phenotype information and variant calls based on whole exome sequencing using the GRCh37 assembly for the Alzheimer’s Disease Sequencing Project (ADSP) Distribution were downloaded from dbGaP (phs000572.v7.p4.c1-c6). Ethical approval and informed consent had been obtained by the researchers who generated this dataset. As described previously, participants were at least 60 years old and were mostly of European-American ancestry though with a small number of Caribbean Hispanic ancestry and all cases met NINCDS-ADRDA criteria for possible, probable or definite AD based on clinical assessment, or had presence of AD (moderate or high likelihood) upon neuropathology examination (Beecham et al., 2017). The cases were chosen whose AD risk score suggested that their disease was not well explained by age, sex or APOE status. Whole exome sequencing was carried out using standard methods as described previously (Bis et al., 2018) and on the ADSP website: https://www.niagads.org/adsp/content/sequencing-pipelines. The downloaded VCF files for autosomal single nucleotide variants and indels were merged into a single file and the PrevAD field was used to define phenotype, yielding 4,600 cases and 6,199 controls.

To obtain population principal components, version 1.90beta of *plink* (https://www.cog-genomics.org/plink2) was run on the genotypes with the options *--maf 0.1 --pca header tabs --make-rel* (Chang et al., 2015; Purcell et al., 2007, 2009).

Each variant was annotated using VEP, PolyPhen and SIFT (Adzhubei et al., 2013; Kumar et al., 2009; McLaren et al., 2016). The same analytic process has previously been applied to an exome sequenced sample of schizophrenia cases and controls (Curtis et al., 2018). The method uses a weighted burden analysis to test whether, in a particular gene or set of genes, variants which are rarer and/or predicted to have more severe functional effects occur more commonly in cases than controls. Variants were weighted according to their functional annotation using the default weights provided with the GENEVARASSOC program, which was used to generate input files for weighted burden analysis by SCOREASSOC (Curtis, 2016, 2012). For example, a weight of 5 was assigned for a synonymous variant, 10 for a non-synonymous variant and 20 for a stop gained variant. Additionally 10 was added to the weight if the PolyPhen annotation was possibly or probably damaging and also if the SIFT annotation was deleterious, meaning that a missense variant annotated as both damaging and deleterious would be assigned an overall weight of 30. The full set of weights is shown in Supplementary Table 1, copied from the previous report (Curtis et al., 2018). Since we reasoned that the effects of any common variants would have been well characterised by previous studies we excluded variants with MAF>0.01 in either cases or controls. SCOREASSOC weights rare variants more strongly than common ones but restricting attention to rare variants meant that this frequency-based weighting had little effect in practice. Variants were also excluded if they did not have a PASS in the information field or if there were more than 10% of genotypes missing in either cases or controls or if the heterozygote count was smaller than both homozygote counts in both cohorts. For each subject a gene-wise weighted burden score is derived as the sum of the variant-wise weights, each multiplied by the number of alleles of the variant which the given subject possesses. If a subject is not genotyped for a variant then they are assigned the subject-wise average score for that variant.

As described previously, ridge regression analysis with lamda=1 was used to test whether the gene-wise score was associated with caseness (Curtis et al., 2018). To do this, SCOREASSOC first calculates the likelihood for the phenotypes as predicted by the first 20 principal components and then calculates the likelihood using a model which additionally incorporates the gene-wise scores. It then carries out a likelihood ratio test assuming that twice the natural log of the likelihood ratio follows a chi-squared distribution with one degree of freedom to produce a p value. Results are expressed as signed log p (SLP) which is the logarithm base 10 of this p value and is positive if the scores are higher in cases and negative if they are higher in controls.

In an attempt to better understand the likely impact of variants on protein structure and function we used visualisation software as described previously (Tsavou and Curtis, 2019). For genes of interest, we considered non-synonymous variants with a weight of at least 50 and visualised the distribution of the altered amino acid residues across functional domains of the protein using the ProteinPaint web application provided at https://pecan.stjude.cloud/home (Zhou et al., 2016). In order to visualise the distribution of altered amino acid residues in the 3D structure of the protein we obtained structures from the RCSB Protein Data Bank at www.rcsb.org (Berman et al., 2000) and colour-coded altered residues using the JSmol viewer provided there and PyMOL 2.4 (“The PyMOL Molecular Graphics System, Version 2.4,” n.d.).

As in the schizophrenia study, pathway analysis was performed using PATHWAYASSOC, which applies the same weighted burden analysis to all the variants seen within a set of genes rather than a single gene, which is easily done by simply summing the gene-wise scores for each subject for all the genes in the set (Curtis, 2016). The same likelihood ratio test, incorporating the first 20 principal components as covariates, was used to test whether the total scores for a set of genes were higher in cases or controls. This was carried out for the 1454 “all GO gene sets, gene symbols” pathways as listed in the file *c5.all.v5.0.symbols.gmt* downloaded from the Molecular Signatures Database at http://www.broadinstitute.org/gsea/msigdb/collections.jsp (Subramanian et al., 2005).

## Results

The weighted burden tests evaluated 1,217,860 variants in 15,777 autosomal genes. Figure 1 shows the QQ plot of the observed SLPs against the distribution expected under the null hypothesis. It can be seen that for almost all genes the results conform well to the SLP=eSLP line. There is one marked outlier, consisting of *TREM2* with an SLP of 14.07, and a few other genes with more extremely positive and negative SLPs than would be expected under the null hypothesis. Ignoring the 200 genes with the highest SLP, the gradient for the genes with positive SLPs was 1.04, with an intercept of −0.016, indicating a small degree of inflation of the test statistic. Ignoring the 200 genes with the lowest SLP, the gradient for the genes with negative SLPs was 1.15, with an intercept of −0.014, indicating a moderate degree of inflation of the test statistic. Table 1 shows results for all genes with an absolute value for the SLP of 3 (equivalent to p=0.001) or greater. 31 genes meet this criterion, while one would expect 16 by chance. Thus, although most do not reach formal standards of statistical significance following correction for multiple testing, some of those listed may represent genes in which variants can affect susceptibility to LOAD and hence seemed worthy of closer examination.

**Figure 1.**
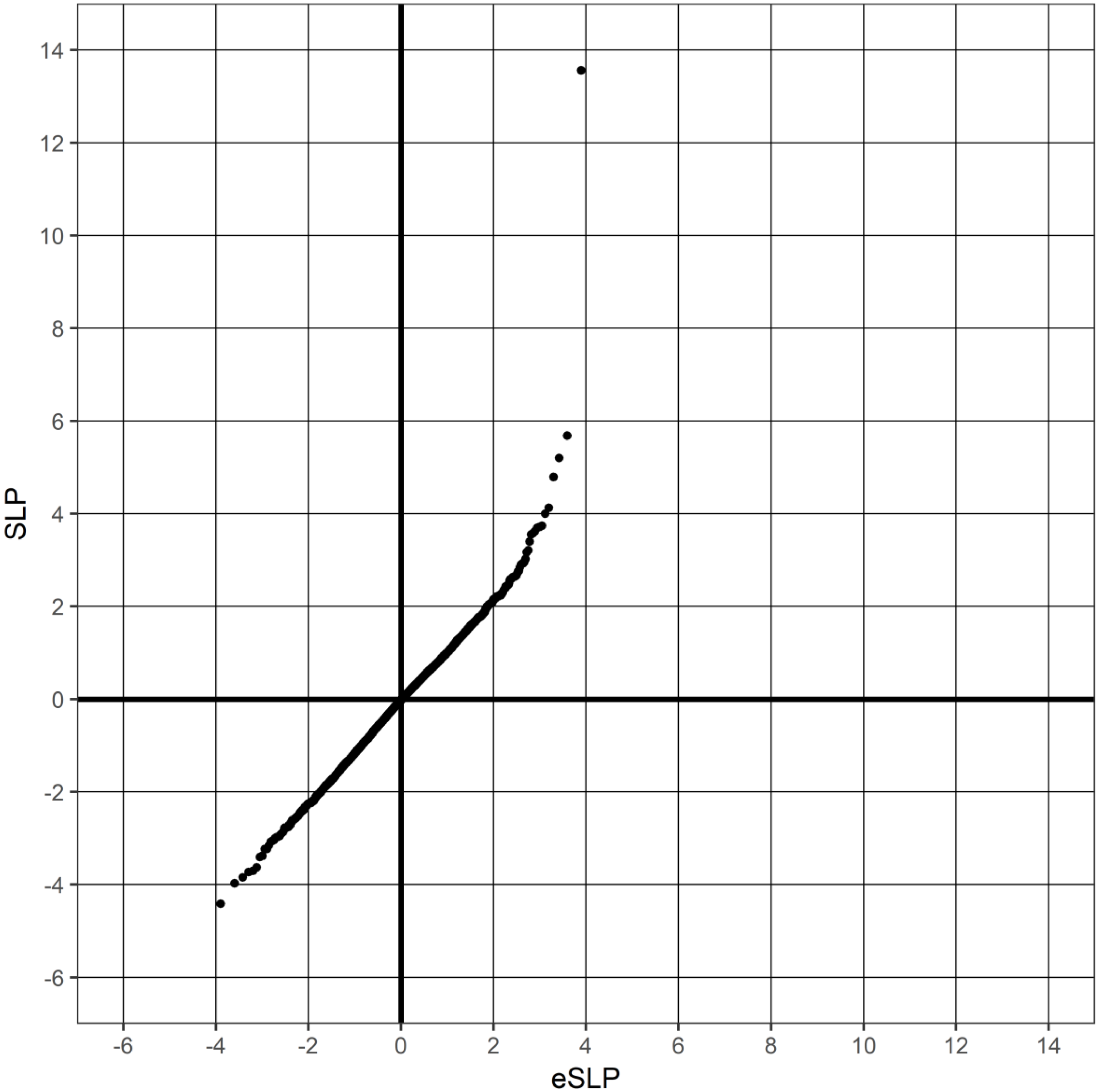
QQ plot of observed versus expected gene-wise SLP.

**Table 1.** List of genes with absolute value for SLP of 3 (equivalent to p = 0.001) or more.

| Gene symbol | SLP | Gene name |
| --- | --- | --- |
| *TREM2* | 13.56 | Triggering receptor expressed on myeloid cells 2 |
| *ABCA7* | 5.68 | ATP binding cassette subfamily A member 7 |
| *OXCT2* | 5.20 | 3-oxoacid CoA-transferase 2 |
| *SORL1* | 4.79 | Sortilin related receptor 1 |
| *WNT7A* | 4.13 | Wnt family member 7A |
| *PIK3R1* | 4.00 | Phosphoinositide-3-kinase regulatory subunit 1 |
| *PSEN1* | 3.74 | Presenilin 1 |
| *EXOC5* | 3.71 | Exocyst complex component 5 |
| *CCDC87* | 3.69 | Coiled-coil domain containing 87 |
| *KLHL35* | 3.62 | Kelch like family member 35 |
| *BMP8B* | 3.58 | Bone Morphogenetic Protein 8b |
| *PRR5-ARHGAP8* | 3.55 | Proline Rich 5 and Rho GTPase Activating Protein 8 |
| *OSBPL8* | 3.40 | Oxysterol binding protein like 8 |
| *C1R* | 3.21 | Complement component 1, R subcomponent |
| *LOC388813* | 3.17 | Uncharacterized Protein ENSP00000383407-Like |
| *H19* | 3.02 | H19 Imprinted Maternally Expressed Transcript |
| *FER1L6-AS1* | -3.00 | FER1L6 Antisense RNA 1 |
| *NDRG2* | -3.04 | NDRG Family Member 2 |
| *TIAF1* | -3.06 | TGFB1-induced anti-apoptotic factor 1 |
| *S100PBP* | -3.08 | S100P binding protein |
| *GFRAL* | -3.15 | GDNF family receptor alpha like |
| *PLA2G12A* | -3.23 | Phospholipase A2 group XIIA |
| *STAC* | -3.23 | SH3 and cysteine rich domain |
| *CBX3* | -3.38 | Chromobox 3 |
| *TCERG1L* | -3.41 | Transcription elongation regulator 1 Like |
| *GADD45G* | -3.63 | Growth arrest and DNA damage inducible gamma |
| *BPIFB6* | -3.70 | BPI fold containing family B member 6 |
| *AIPL1* | -3.73 | Aryl hydrocarbon receptor interacting protein like 1 |
| *TBC1D23* | -3.84 | TBC1 domain family member 23 |
| *PISD* | -3.97 | Phosphatidylserine decarboxylase |
| *MUS81* | -4.41 | MUS81 structure-specific endonuclease subunit |

Of genes with SLPs above 3, a number have previously been implicated by sequencing studies, consisting of *TREM2, ABCA7, SORL1* and *PSEN1* (Blue et al., 2018; Guerreiro et al., 2013; Kim, 2018; Patel et al., 2019; Raghavan et al., 2018). Many of the others did not seem likely to have any plausible role in AD susceptibility but there were some exceptions which we list here. *PIK3R1* codes for a key component of the PI3K/Akt/GSK-3β signalling pathway and in neuronally differentiated PC12 cells activation of this pathway is neuroprotective against Aβ_25-35_-induced apoptosis and tau hyperphosphorylation (Cheng et al., 2018). Conversely, in a mouse model of AD Aβ oligomers block activation of this pathway, leading to decreased neuronal survival (Jimenez et al., 2011). TMP21 dysregulation is implicated in AD pathogenesis and in TMP21 knock-down mice *PiK3r1* was significantly down-regulated and was one of the top eight dysregulated genes in both the hippocampus and cortex gene signal networks, in which it was a hub gene (Zhang et al., 2019). *WNT7A* is expressed in hippocampal neurons and in mouse models Wnt signalling dysfunction leads to production and aggregation of Aβ, tau phosphorylation and hippocampus-dependent cognitive impairment (Tapia-Rojas and Inestrosa, 2018). It has been reported that the R subcomponent of complement component 1, the product of *C1R*, is enriched in amyloid plaques from patients with Alzheimer’s disease compared to amyloid plaques from subjects without (Xiong et al., 2019). Heterozygous missense variants in *C1R* are a known cause of periodontal Ehlers-Danlos syndrome (pEDS) (Kapferer-Seebacher et al., 2016) and this is potentially of interest for two reasons. One is that a recent MRI study of seven adults with pEDS and *C1R* mutations reported that they all had leukoencephalopathy, which appeared to be progressive with age and which was suggestive of underlying small vessel disease (Kapferer-Seebacher et al., 2019). The second is that pEDS is characterised by periodontitis and it has been proposed that periodontal disease is a risk factor for LOAD, perhaps mediated either by direct infection of brain with the key periodontal pathogen *Porphyromonas gingivalis* or through secondary effects of chronic inflammation, perhaps involving activation of the complement cascade (Singhrao et al., 2015). Although there is no direct evidence to implicate *EXOC5* in LOAD, in a mouse model cell-specific deletion of *Exoc5* results in progressive hearing loss associated with loss of hair cells and spiral ganglion neurons (Lee et al., 2018).

With regard to the genes with SLP less than −3 shown in Table 1, the fact that *TIAF1* produces an SLP of −3.06 is of some interest because of a report that its product can self-aggregate and that polymerised TIAF1 physically interacts with amyloid fibrils, putatively promoting plaque formation (Lee et al., 2010). *NDRG2* expression is increased at the protein and mRNA level in brains of subjects with AD and the protein is detectable in both dystrophic neurons and amyloid plaques (Mitchelmore et al., 2004). *NDRG2* is up-regulated in mouse models of senescence and AD and in cellular models knock-down of *NDRG2* reduces BACE1 and Aβ1-42 levels while NDRG2 overexpression increases them (Rong et al., 2017). *SOX14* codes for a transcription factor which is required for normal neuronal development in the cerebellum but there seems to be little evidence for its involvement in LOAD (Prekop et al., 2018).

The results for other genes previously implicated in AD susceptibility were examined. There was no difference in the weighted burden scores between cases and controls for *APP* (SLP=0.11) and there were no valid variants for *PSEN2* or *APOE*.

In general large numbers of variants which were very rare contributed to the observed SLPs in the genes listed above and as each variant might occur in only one or two subjects one could not make any firm inference about their individual effect on risk. The product of transcript NM_00002.1 of *PSEN1* consists of 468 amino acids but 28 out of 33 of the highly weighted variants (weight>=50) in cases affected residues between positions 35 and 270 and this also applied to 14 out of 17 highly weighted variants seen in controls. That is, the highly weighted variants in general tended to affect this region and overall there were more of them among cases than controls. Visualisation suggested that these amino acids possibly tended to occur in the regions involved in APP recognition as observed from the cryo-EM structure of the cross-linked human γ-secretase–APP-C83 complex (Zhou et al., 2019). The mature product of the NM_001733 transcript of *C1R* consists of 628 amino acids and in a series of pedigrees with pEDS 14 out of 15 segregated a variant affecting a residue at a position between 272 and 417 (Kapferer-Seebacher et al., 2016). Only 19 out of 58 highly weighted variants in cases affected amino acids in this region compared with 14 out of 41 in controls and none of these amino acids corresponded with any of the ones affected in the pEDS pedigrees. Thus, there was little suggestion that the region of the protein implicated in pEDS was disproportionately affected in the LOAD cases. In fact, affected residues in cases tended to cluster in the region between positions 430 and 537, which accounted for 17 out of 58 in cases and only 2 out of 48 in controls. However the small number of counts for the individual variants precludes drawing any firm conclusions and in other genes, visualisation of the distribution of altered amino acids did not reveal any obvious patterns. Table 2 shows the genotype counts for variants in all the above-named genes for those variants with a minor allele count of at least 10 and an odds ratio over 2, or, for the genes with SLP<-3, less than 0.5. The variants in *TREM2* have been reported in a previous analysis of this dataset (Song et al., 2017). Likewise, deleterious mutations in *ABCA7* are known to be risk factors for both early and late onset Alzheimer’s disease (De Roeck et al., 2017). The variants in *SORL1* at 11:121348842 (rs140888526), 11:121421313 (rs148430425) and 11:121421361 (rs146438170) have not previously been implicated in LOAD susceptibility and although the gene-based SLP of 4.42 suggests that overall there is an excess of functionally significant variants among cases the numbers are too small to draw any definitive conclusions about any of these variants individually. The three qualifying variants listed for *C1R* are not predicted to impact on protein function. The variant in *EXOC5* at 14:57714397 (rs199912312) is predicted to be deleterious and probably damaging and is seen in 14 cases and only 3 controls but has not been reported to be associated with any phenotype. The variant in *TIAF1* at 17:27401061 (rs73986791) is not predicted to affect protein function and has not previously been reported.

**Table 2.**
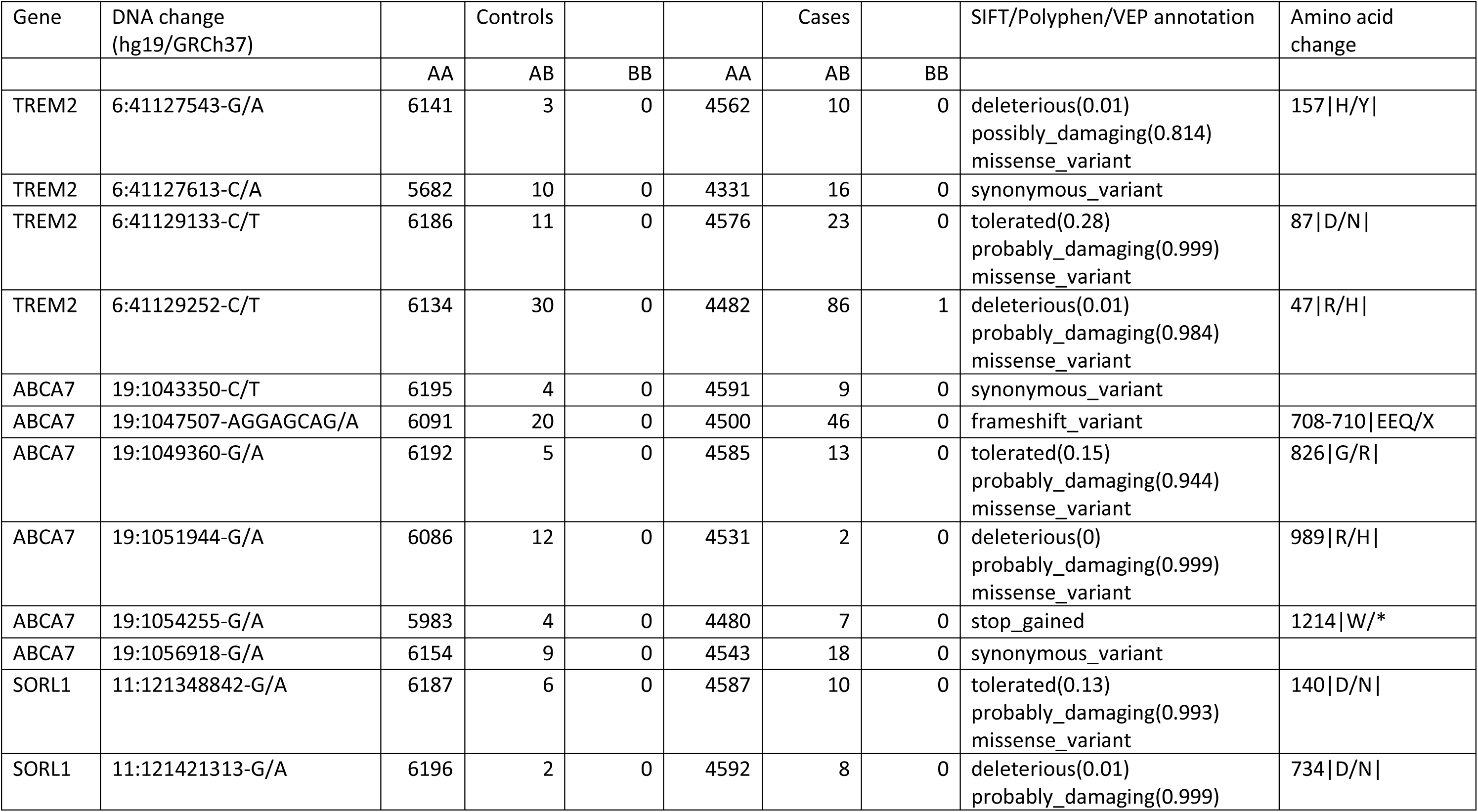
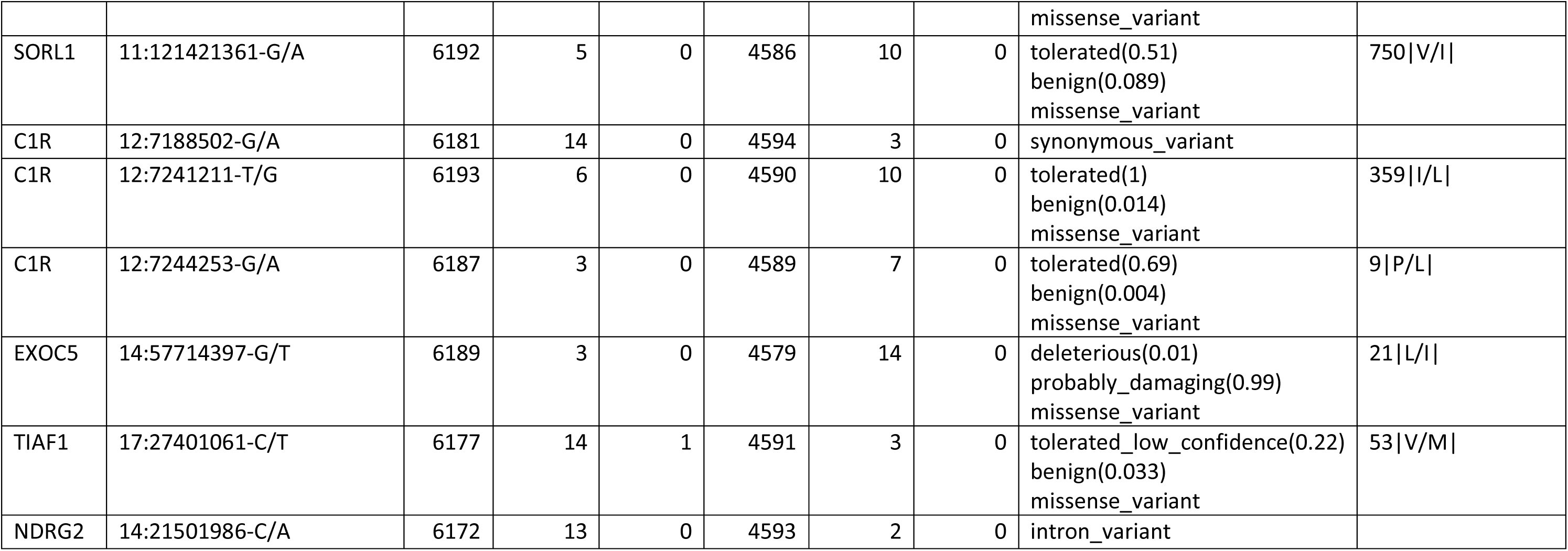
List of variants in genes of interest with total minor allele count of at least 10 and odds ratio greater than 2 or, for genes with negative SLP, less than 0.5.

In the gene set analysis using PATHWAYASSOC the highest SLP achieved was 3.11 for the PROTEIN_MATURATION set, which given that 1454 sets were tested is within chance expectation. Examination of the output suggested that this was driven largely by the inclusion of *PSEN1* in this set and no other gene in it had an individual SLP higher than 1.3 (p=0.05). Table 3 lists the gene sets with SLP of −3 or less. The most negative is PROTEIN_TYROSINE_PHOSPHATASE_ACTIVITY with SLP=-5.27. This result is of interest because it indicates that controls are more likely to carry functionally significant variants than cases, suggesting that impairment of this activity might be protective against LOAD. Tyrosine phosphatases counter the activities of tyrosine kinases and for example the loss of *PTPRS*, which has an SLP of −2.41, has been shown to cause activation of the EGFR/PI3K pathway (Morris et al., 2011). There is an extensive literature on the role of tyrosine phosphatase in Alzheimer pathology and genetic reduction of striatal-enriched tyrosine phosphatase has been shown to reverse cognitive and cellular deficits in mouse models of AD (Hendriks et al., 2013; Zhang et al., 2010). A recent study has shown that bergenin, a tyrosine phosphatase inhibitor, has neuroprotective effects in a rat model of AD (Barai et al., 2019). There are 47 genes within this set and Table 4 lists those which individually have SLP of −1.3 (equivalent to p=0.05) or less. It can be seen that the genes for protein tyrosine phosphatase, receptor types R, S, T and U all have SLPs of approximately −2. *PTPN1* has SLP of −1.31 (p=0.05) and treatment with a non-specific inhibitor of its product, PTP1B, using sodium orthovanadate has recently been reported to reduce cognitive symptoms in a mouse model of vascular dementia (Kumar et al., 2019). Another study in animal models showing that activation of the PI3K/Akt pathway combined with inhibition of PTP1B suppresses Aβ-induced GSK-3β activation and tau phosphorylation (Kanno et al., 2016).

**Table 3.** List of gene sets achieving SLP of −3 (equivalent to p = 0.001) or less.

| Gene set | SLP |
| --- | --- |
| PROTEIN_TYROSINE_PHOSPHATASE_ACTIVITY | -5.27 |
| PHOSPHOPROTEIN_PHOSPHATASE_ACTIVITY | -4.50 |
| PHOSPHORIC_MONOESTER_HYDROLASE_ACTIVITY | -3.86 |
| TRANSMEMBRANE_RECEPTOR_ACTIVITY | -3.75 |
| NUCLEUS | -3.63 |
| TRANSMEMBRANE_RECEPTOR_PROTEIN_PHOSPHATASE_ACTIVITY | -3.53 |

**Table 4.** Genes in the set PROTEIN_TYROSINE_PHOSPHATASE_ACTIVITY with SLPs of −1.3 (equivalent to p=0.05) or less.

| Gene symbol | SLP | Gene name |
| --- | --- | --- |
| PTPRS | -2.41 | Protein tyrosine phosphatase, receptor type S |
| PTPRU | -2.30 | Protein tyrosine phosphatase, receptor type U |
| PTPRR | -2.06 | Protein tyrosine phosphatase, receptor type R |
| DUSP6 | -2.05 | Dual specificity phosphatase 6 |
| PTPRT | -1.96 | Protein tyrosine phosphatase, receptor type T |
| PTPN12 | -1.66 | Protein tyrosine phosphatase, non-receptor type 12 |
| PTP4A3 | -1.32 | Protein tyrosine phosphatase type IVA, member 3 |
| PTPN22 | -1.30 | Protein tyrosine phosphatase, non-receptor type 22 |
| PTPN1 | -1.30 | Protein tyrosine phosphatase, non-receptor type 1 |

Most of the other gene sets which produced strongly negative SLPs did so because they also contained tyrosine phosphatase genes. For those which did not, inspection of the outputs did not reveal any genes which seemed likely to be related to AD susceptibility.

There have been reports that genetic reduction or inhibition of a different tyrosine phosphatase, striatum-enriched protein-tyrosine phosphatase (STEP), reverses cognitive and cellular deficits in a mouse model of Alzheimer’s disease (Xu et al., 2014; Zhang et al., 2010). STEP is coded for by *PTPN5* and there was no significant evidence for an increased variant burden in this gene in controls (SLP=-0.69, NS).

## Discussion

*TREM2, ABCA7* and *SORL1*, the genes with the first, second and fourth highest SLPs, are well-established susceptibility genes for LOAD, which inspires some confidence in the validity of the approach. Although a previous analysis of this dataset did find some support for the involvement of *PSEN1*, only specific variants were included and the results were significant only at p<0.05. The current analysis includes all rare variants and yields an SLP of 3.74, equivalent to p=0.0002. This provides stronger evidence and also suggests that a wider range of *PSEN1* variants may have an effect on LOAD risk. The results for *PIK3R1*, W*NT7A* and *C1R*, considered alongside the relevant literature, suggest that variants affecting functioning of these genes may increase LOAD susceptibility and for each of these genes there are previous reports that could be interpreted as lending further support to this hypothesis but it is not possible at present to make definitive claims. It is unclear whether the results for *EXOC5* reflect any influence on LOAD susceptibility or whether they have occurred simply by chance. Aside from the risk variants previously reported for *TREM2* and *ABCA7*, the current study is unable to implicate specific variants rather than genes.

Also of interest are the results for genes in which the SLP is strongly negative, implying that variants may protect against the development of LOAD. Perhaps the most plausible result is for *TIAF1* because of the claim that TIAF1 self-aggregates and leads to apoptosis, Aβ formation and plaque formation (Lee et al., 2010). If this is so then variants which interfered with this process would be expected to reduce LOAD risk. Likewise, there is some prior evidence that *NDRG2* could be involved in aspects of LOAD pathogenesis and the results we obtain suggest that variants in it might be protective but additional studies would be needed to settle this question.

Probably the most striking result is the evidence for a protective effect from variants in the set of genes encoding tyrosine phosphatases, which produces an SLP of −5.27 (equivalent to p = 5 × 10^−6^). Given the established role of tyrosine phosphatase processes in the development of AD this result is eminently plausible but has not been described previously. Conversely, the results for *PIK3R1*, which has a reciprocal effect, suggest that variants in this gene can increase risk of LOAD. Taken together, these findings suggest that increased activity of the PI3K/Akt signalling pathway is protective against LOAD whereas reduced activity increases risk, in accordance with the study of PI3K/Akt activation and PTP1B inhibition referred to previously (Kanno et al., 2016).

Analysing all variants together in a single joint analysis makes good use of the available data while avoiding problems due to multiple testing issues. The results obtained serve to highlight genes rather than individual variants, which tend to be too rare for individual inferences to be made. The results reported here serve to reinforce some known aspects of LOAD biology and to suggest additional leads which might profitably be pursued.

## Acknowledgments

The author wishes to acknowledge the staff supporting the High Performance Computing Cluster, Computer Science Department, University College London. This work was carried out in part using resources provided by BBSRC equipment grant BB/R01356X/1.

The Alzheimer’s Disease Sequencing Project (ADSP) is comprised of two Alzheimer’s Disease (AD) genetics consortia and three National Human Genome Research Institute (NHGRI) funded Large Scale Sequencing and Analysis Centers (LSAC). The two AD genetics consortia are the Alzheimer’s Disease Genetics Consortium (ADGC) funded by NIA (U01 AG032984), and the Cohorts for Heart and Aging Research in Genomic Epidemiology (CHARGE) funded by NIA (R01 AG033193), the National Heart, Lung, and Blood Institute (NHLBI), other National Institute of Health (NIH) institutes and other foreign governmental and non-governmental organizations. The Discovery Phase analysis of sequence data is supported through UF1AG047133 (to Drs. Schellenberg, Farrer, Pericak-Vance, Mayeux, and Haines); U01AG049505 to Dr. Seshadri; U01AG049506 to Dr. Boerwinkle; U01AG049507 to Dr. Wijsman; and U01AG049508 to Dr. Goate and the Discovery Extension Phase analysis is supported through U01AG052411 to Dr. Goate, U01AG052410 to Dr. Pericak-Vance and U01 AG052409 to Drs. Seshadri and Fornage. Data generation and harmonization in the Follow-up Phases is supported by U54AG052427 (to Drs. Schellenberg and Wang).

The ADGC cohorts include: Adult Changes in Thought (ACT), the Alzheimer’s Disease Centers (ADC), the Chicago Health and Aging Project (CHAP), the Memory and Aging Project (MAP), Mayo Clinic (MAYO), Mayo Parkinson’s Disease controls, University of Miami, the Multi-Institutional Research in Alzheimer’s Genetic Epidemiology Study (MIRAGE), the National Cell Repository for Alzheimer’s Disease (NCRAD), the National Institute on Aging Late Onset Alzheimer’s Disease Family Study (NIA-LOAD), the Religious Orders Study (ROS), the Texas Alzheimer’s Research and Care Consortium (TARC), Vanderbilt University/Case Western Reserve University (VAN/CWRU), the Washington Heights-Inwood Columbia Aging Project (WHICAP) and the Washington University Sequencing Project (WUSP), the Columbia University Hispanic-Estudio Familiar de Influencia Genetica de Alzheimer (EFIGA), the University of Toronto (UT), and Genetic Differences (GD).

The CHARGE cohorts are supported in part by National Heart, Lung, and Blood Institute (NHLBI) infrastructure grant HL105756 (Psaty), RC2HL102419 (Boerwinkle) and the neurology working group is supported by the National Institute on Aging (NIA) R01 grant AG033193. The CHARGE cohorts participating in the ADSP include the following: Austrian Stroke Prevention Study (ASPS), ASPS-Family study, and the Prospective Dementia Registry-Austria (ASPS/PRODEM-Aus), the Atherosclerosis Risk in Communities (ARIC) Study, the Cardiovascular Health Study (CHS), the Erasmus Rucphen Family Study (ERF), the Framingham Heart Study (FHS), and the Rotterdam Study (RS). ASPS is funded by the Austrian Science Fond (FWF) grant number P20545-P05 and P13180 and the Medical University of Graz. The ASPS-Fam is funded by the Austrian Science Fund (FWF) project I904),the EU Joint Programme - Neurodegenerative Disease Research (JPND) in frame of the BRIDGET project (Austria, Ministry of Science) and the Medical University of Graz and the Steiermärkische Krankenanstalten Gesellschaft. PRODEM-Austria is supported by the Austrian Research Promotion agency (FFG) (Project No. 827462) and by the Austrian National Bank (Anniversary Fund, project 15435. ARIC research is carried out as a collaborative study supported by NHLBI contracts (HHSN268201100005C, HHSN268201100006C, HHSN268201100007C, HHSN268201100008C, HHSN268201100009C, HHSN268201100010C, HHSN268201100011C, and HHSN268201100012C). Neurocognitive data in ARIC is collected by U01 2U01HL096812, 2U01HL096814, 2U01HL096899, 2U01HL096902, 2U01HL096917 from the NIH (NHLBI, NINDS, NIA and NIDCD), and with previous brain MRI examinations funded by R01-HL70825 from the NHLBI. CHS research was supported by contracts HHSN268201200036C, HHSN268200800007C, N01HC55222, N01HC85079, N01HC85080, N01HC85081, N01HC85082, N01HC85083, N01HC85086, and grants U01HL080295 and U01HL130114 from the NHLBI with additional contribution from the National Institute of Neurological Disorders and Stroke (NINDS). Additional support was provided by R01AG023629, R01AG15928, and R01AG20098 from the NIA. FHS research is supported by NHLBI contracts N01-HC-25195 and HHSN268201500001I. This study was also supported by additional grants from the NIA (R01s AG054076, AG049607 and AG033040 and NINDS (R01 NS017950). The ERF study as a part of EUROSPAN (European Special Populations Research Network) was supported by European Commission FP6 STRP grant number 018947 (LSHG-CT-2006-01947) and also received funding from the European Community’s Seventh Framework Programme (FP7/2007-2013)/grant agreement HEALTH-F4-2007-201413 by the European Commission under the programme “Quality of Life and Management of the Living Resources” of 5th Framework Programme (no. QLG2-CT-2002-01254). High-throughput analysis of the ERF data was supported by a joint grant from the Netherlands Organization for Scientific Research and the Russian Foundation for Basic Research (NWO-RFBR 047.017.043). The Rotterdam Study is funded by Erasmus Medical Center and Erasmus University, Rotterdam, the Netherlands Organization for Health Research and Development (ZonMw), the Research Institute for Diseases in the Elderly (RIDE), the Ministry of Education, Culture and Science, the Ministry for Health, Welfare and Sports, the European Commission (DG XII), and the municipality of Rotterdam. Genetic data sets are also supported by the Netherlands Organization of Scientific Research NWO Investments (175.010.2005.011, 911-03-012), the Genetic Laboratory of the Department of Internal Medicine, Erasmus MC, the Research Institute for Diseases in the Elderly (014-93-015; RIDE2), and the Netherlands Genomics Initiative (NGI)/Netherlands Organization for Scientific Research (NWO) Netherlands Consortium for Healthy Aging (NCHA), project 050-060-810. All studies are grateful to their participants, faculty and staff. The content of these manuscripts is solely the responsibility of the authors and does not necessarily represent the official views of the National Institutes of Health or the U.S. Department of Health and Human Services.

The four LSACs are: the Human Genome Sequencing Center at the Baylor College of Medicine (U54 HG003273), the Broad Institute Genome Center (U54HG003067), The American Genome Center at the Uniformed Services University of the Health Sciences (U01AG057659), and the Washington University Genome Institute (U54HG003079).

Biological samples and associated phenotypic data used in primary data analyses were stored at Study Investigators institutions, and at the National Cell Repository for Alzheimer’s Disease (NCRAD, U24AG021886) at Indiana University funded by NIA. Associated Phenotypic Data used in primary and secondary data analyses were provided by Study Investigators, the NIA funded Alzheimer’s Disease Centers (ADCs), and the National Alzheimer’s Coordinating Center (NACC, U01AG016976) and the National Institute on Aging Genetics of Alzheimer’s Disease Data Storage Site (NIAGADS, U24AG041689) at the University of Pennsylvania, funded by NIA, and at the Database for Genotypes and Phenotypes (dbGaP) funded by NIH. This research was supported in part by the Intramural Research Program of the National Institutes of health, National Library of Medicine. Contributors to the Genetic Analysis Data included Study Investigators on projects that were individually funded by NIA, and other NIH institutes, and by private U.S. organizations, or foreign governmental or nongovernmental organizations.

## Data availability statement

The data used was downloaded from dbGaP so data sharing is not applicable to this article as no new data was created in this study. However scripts, supporting files, interim files and full results have been deposited at the NIAGADS site: https://www.niagads.org/datasets/ng00091.

---

**Supplementary table S1.** The table shows the weight accorded to each type of variant as annotated by VEP (McLaren et al., 2016). 10 was added to this weight if the variant was annotated by Polyphen as possibly or probably damaging and 10 was added if SIFT annotated it as deleterious (Adzhubei et al., 2013; Kumar et al., 2009).

| VEP annotation | Weight |
| --- | --- |
| intergenic_variant | 1 |
| feature_truncation | 3 |
| regulatory_region_variant | 3 |
| feature_elongation | 3 |
| regulatory_region_amplification | 3 |
| regulatory_region_ablation | 3 |
| TF_binding_site_variant | 3 |
| TFBS_amplification | 3 |
| TFBS_ablation | 3 |
| downstream_gene_variant | 3 |
| upstream_gene_variant | 3 |
| non_coding_transcript_variant | 3 |
| NMD_transcript_variant | 3 |
| intron_variant | 3 |
| non_coding_transcript_exon_variant | 3 |
| 3_prime_UTR_variant | 5 |
| 5_prime_UTR_variant | 5 |
| mature_miRNA_variant | 5 |
| coding_sequence_variant | 5 |
| synonymous_variant | 5 |
| stop_retained_variant | 5 |
| incomplete_terminal_codon_variant | 5 |
| splice_region_variant | 5 |
| protein_altering_variant | 10 |
| missense_variant | 10 |
| inframe_deletion | 15 |
| inframe_insertion | 15 |
| transcript_amplification | 15 |
| start_lost | 15 |
| stop_lost | 15 |
| frameshift_variant | 20 |
| stop_gained | 20 |
| splice_donor_variant | 20 |
| splice_acceptor_variant | 20 |
| transcript_ablation | 20 |

## References

Adzhubei, I., Jordan, D.M., Sunyaev, S.R. (2013) Predicting functional effect of human missense mutations using PolyPhen-2. Curr. Protoc. Hum. Genet. 7 Unit7.20.

Barai, P., Raval, N., Acharya, S., Borisa, A., Bhatt, H., Acharya, N. (2019) Neuroprotective effects of bergenin in Alzheimer’s disease: Investigation through molecular docking, in vitro and in vivo studies. Behav. Brain Res. 356, 18–40.

Beecham, G.W., Bis, J.C., Martin, E.R., Choi, S.-H., DeStefano, A.L., van Duijn, C.M., Fornage, M., Gabriel, S.B., Koboldt, D.C., Larson, D.E., Naj, A.C., Psaty, B.M., Salerno, W., Bush, W.S., Foroud, T.M., Wijsman, E., Farrer, L.A., Goate, A., Haines, J.L., Pericak-Vance, M.A., Boerwinkle, E., Mayeux, R., Seshadri, S., Schellenberg, G. (2017) The Alzheimer’s Disease Sequencing Project: Study design and sample selection. Neurol. Genet. 3, e194.

Berman, H.M., Westbrook, J., Feng, Z., Gilliland, G., Bhat, T.N., Weissig, H., Shindyalov, I.N., Bourne, P.E. (2000) The Protein Data Bank. Nucleic Acids Res. 28, 235–42.

Bis, J.C., Jian, X., Kunkle, B.W., Chen, Y., Hamilton-Nelson, K.L., Bush, W.S., Salerno, W.J., Lancour, D., Ma, Y., Renton, A.E., Marcora, E., Farrell, J.J., Zhao, Y., Qu, L., Ahmad, S., Amin, N., Amouyel, P., Beecham, G.W., Below, J.E., Campion, D., Charbonnier, C., Chung, J., Crane, P.K., Cruchaga, C., Cupples, L.A., Dartigues, J.-F., Debette, S., Deleuze, J.-F., Fulton, L., Gabriel, S.B., Genin, E., Gibbs, R.A., Goate, A., Grenier-Boley, B., Gupta, N., Haines, J.L., Havulinna, A.S., Helisalmi, S., Hiltunen, M., Howrigan, D.P., Ikram, M.A., Kaprio, J., Konrad, J., Kuzma, A., Lander, E.S., Lathrop, M., Lehtimäki, T., Lin, H., Mattila, K., Mayeux, R., Muzny, D.M., Nasser, W., Neale, B., Nho, K., Nicolas, G., Patel, D., Pericak-Vance, M.A., Perola, M., Psaty, B.M., Quenez, O., Rajabli, F., Redon, R., Reitz, C., Remes, A.M., Salomaa, V., Sarnowski, C., Schmidt, H., Schmidt, M., Schmidt, R., Soininen, H., Thornton, T.A., Tosto, G., Tzourio, C., van der Lee, S.J., van Duijn, C.M., Vardarajan, B., Wang, W., Wijsman, E., Wilson, R.K., Witten, D., Worley, K.C., Zhang, X., Bellenguez, C., Lambert, J.-C., Kurki, M.I., Palotie, A., Daly, M., Boerwinkle, E., Lunetta, K.L., Destefano, A.L., Dupuis, J., Martin, E.R., Schellenberg, G.D., Seshadri, S., Naj, A.C., Fornage, M., Farrer, L.A. (2018) Whole exome sequencing study identifies novel rare and common Alzheimer’s-Associated variants involved in immune response and transcriptional regulation. Mol. Psychiatry 1.

Blue, E.E., Bis, J.C., Dorschner, M.O., Tsuang, D.W., Barral, S.M., Beecham, G., Below, J.E., Bush, W.S., Butkiewicz, M., Cruchaga, C., DeStefano, A., Farrer, L.A., Goate, A., Haines, J., Jaworski, J., Jun, G., Kunkle, B., Kuzma, A., Lee, J.J., Lunetta, K.L., Ma, Y., Martin, E., Naj, A., Nato, A.Q., Navas, P., Nguyen, H., Reitz, C., Reyes, D., Salerno, W., Schellenberg, G.D., Seshadri, S., Sohi, H., Thornton, T.A., Valadares, O., van Duijn, C., Vardarajan, B.N., Wang, L.-S., Boerwinkle, E., Dupuis, J., Pericak-Vance, M.A., Mayeux, R., Wijsman, E.M., on behalf of the Alzheimer’s Disease Sequencing Project (2018) Genetic Variation in Genes Underlying Diverse Dementias May Explain a Small Proportion of Cases in the Alzheimer’s Disease Sequencing Project. Dement. Geriatr. Cogn. Disord. 45, 1–17.

Chang, C.C., Chow, C.C., Tellier, L.C., Vattikuti, S., Purcell, S.M., Lee, J.J. (2015) Second-generation PLINK: rising to the challenge of larger and richer datasets. Gigascience 4, 7.

Cheng, W., Chen, W., Wang, P., Chu, J. (2018) Asiatic acid protects differentiated PC12 cells from Aβ25–35-induced apoptosis and tau hyperphosphorylation via regulating PI3K/Akt/GSK-3β signaling. Life Sci. 208, 96–101.

Curtis, D. (2012) A rapid method for combined analysis of common and rare variants at the level of a region, gene, or pathway. Adv Appl Bioinform Chem 5, 1–9.

Curtis, D. (2016) Pathway analysis of whole exome sequence data provides further support for the involvement of histone modification in the aetiology of schizophrenia. Psychiatr. Genet. 26, 223–7.

Curtis, D., Coelewij, L., Liu, S.-H., Humphrey, J., Mott, R. (2018) Weighted Burden Analysis of Exome-Sequenced Case-Control Sample Implicates Synaptic Genes in Schizophrenia Aetiology. Behav. Genet. 203521.

Curtis, D., UK10K Consortium (2016) Practical experience of the application of a weighted burden test to whole exome sequence data for obesity and schizophrenia. Ann Hum Genet 80, 38–49.

De Roeck, A., Van den Bossche, T., van der Zee, J., Verheijen, J., De Coster, W., Van Dongen, J., Dillen, L., Baradaran-Heravi, Y., Heeman, B., Sanchez-Valle, R., Lladó, A., Nacmias, B., Sorbi, S., Gelpi, E., Grau-Rivera, O., Gómez-Tortosa, E., Pastor, P., Ortega-Cubero, S., Pastor, M.A., Graff, C., Thonberg, H., Benussi, L., Ghidoni, R., Binetti, G., de Mendonça, A., Martins, M., Borroni, B., Padovani, A., Almeida, M.R., Santana, I., Diehl-Schmid, J., Alexopoulos, P., Clarimon, J., Lleó, A., Fortea, J., Tsolaki, M., Koutroumani, M., Matěj, R., Rohan, Z., De Deyn, P., Engelborghs, S., Cras, P., Van Broeckhoven, C., Sleegers, K., consortium, O. behalf of the E.E.-O.D. (EU E. (2017) Deleterious ABCA7 mutations and transcript rescue mechanisms in early onset Alzheimer’s disease. Acta Neuropathol. 134, 475–487.

Guerreiro, R., Wojtas, A., Bras, J., Carrasquillo, M., Rogaeva, E., Majounie, E., Cruchaga, C., Sassi, C., Kauwe, J.S.K., Younkin, S., Hazrati, L., Collinge, J., Pocock, J., Lashley, T., Williams, J., Lambert, J.-C., Amouyel, P., Goate, A., Rademakers, R., Morgan, K., Powell, J., St. George-Hyslop, P., Singleton, A., Hardy, J. (2013) *TREM2* Variants in Alzheimer’s Disease. N. Engl. J. Med. 368, 117–127.

Hendriks, W.J.A.J., Elson, A., Harroch, S., Pulido, R., Stoker, A., den Hertog, J. (2013) Protein tyrosine phosphatases in health and disease. FEBS J. 280, 708–730.

Jimenez, S., Torres, M., Vizuete, M., Sanchez-Varo, R., Sanchez-Mejias, E., Trujillo-Estrada, L., Carmona-Cuenca, I., Caballero, C., Ruano, D., Gutierrez, A., Vitorica, J. (2011) Age-dependent Accumulation of Soluble Amyloid β (Aβ) Oligomers Reverses the Neuroprotective Effect of Soluble Amyloid Precursor Protein-α (sAPPα) by Modulating Phosphatidylinositol 3-Kinase (PI3K)/Akt-GSK-3β Pathway in Alzheimer Mouse Model. J. Biol. Chem. 286, 18414–18425.

Kanno, T., Tsuchiya, A., Tanaka, A., Nishizaki, T. (2016) Combination of PKCε Activation and PTP1B Inhibition Effectively Suppresses Aβ-Induced GSK-3β Activation and Tau Phosphorylation. Mol. Neurobiol. 53, 4787–4797.

Kapferer-Seebacher, I., Pepin, M., Werner, R., Aitman, T.J., Nordgren, A., Stoiber, H., Thielens, N., Gaboriaud, C., Amberger, A., Schossig, A., Gruber, R., Giunta, C., Bamshad, M., Björck, E., Chen, C., Chitayat, D., Dorschner, M., Schmitt-Egenolf, M., Hale, C.J., Hanna, D., Hennies, H.C., Heiss-Kisielewsky, I., Lindstrand, A., Lundberg, P., Mitchell, A.L., Nickerson, D.A., Reinstein, E., Rohrbach, M., Romani, N., Schmuth, M., Silver, R., Taylan, F., Vandersteen, A., Vandrovcova, J., Weerakkody, R., Yang, M., Pope, F.M., Aleck, K., Banki, Z., Dudas, J., Dumfahrt, H., Haririan, H., Hartsfield, J.K., Kagen, C.N., Lindert, U., Meitinger, T., Posch, W., Pritz, C., Ross, D., Schroer, R.J., Wick, G., Wildin, R., Wilflingseder, D., Byers, P.H., Zschocke, J. (2016) Periodontal Ehlers-Danlos Syndrome Is Caused by Mutations in C1R and C1S, which Encode Subcomponents C1r and C1s of Complement. Am. J. Hum. Genet. 99, 1005–1014.

Kapferer-Seebacher, I., Waisfisz, Q., Boesch, S., Bronk, M., van Tintelen, P., Gizewski, E.R., Groebner, R., Zschocke, J., van der Knaap, M.S. (2019) Periodontal Ehlers–Danlos syndrome is associated with leukoencephalopathy. Neurogenetics 20, 1–8.

Kim, J.H. (2018) Genetics of Alzheimer’s Disease. Dement. Neurocognitive Disord. 17, 131.

Kumar, P., Henikoff, S., Ng, P.C. (2009) Predicting the effects of coding non-synonymous variants on protein function using the SIFT algorithm. Nat. Protoc. 4, 1073–1081.

Kumar, S., Ivanov, S., Lagunin, A., Goel, R.K. (2019) Attenuation of hyperhomocysteinemia induced vascular dementia by sodium orthovanadate perhaps via PTP1B: Pertinent downstream outcomes. Behav. Brain Res. 364, 29–40.

Kunkle, B.W., Grenier-Boley, B., Sims, R., Bis, J.C., Damotte, V., Naj, A.C., Boland, A., Vronskaya, M., van der Lee, S.J., Amlie-Wolf, A., Bellenguez, C., Frizatti, A., Chouraki, V., Martin, E.R., Sleegers, K., Badarinarayan, N., Jakobsdottir, J., Hamilton-Nelson, K.L., Moreno-Grau, S., Olaso, R., Raybould, R., Chen, Y., Kuzma, A.B., Hiltunen, M., Morgan, T., Ahmad, S., Vardarajan, B.N., Epelbaum, J., Hoffmann, P., Boada, M., Beecham, G.W., Garnier, J.-G., Harold, D., Fitzpatrick, A.L., Valladares, O., Moutet, M.-L., Gerrish, A., Smith, A. V., Qu, L., Bacq, D., Denning, N., Jian, X., Zhao, Y., Del Zompo, M., Fox, N.C., Choi, S.-H., Mateo, I., Hughes, J.T., Adams, H.H., Malamon, J., Sanchez-Garcia, F., Patel, Y., Brody, J.A., Dombroski, B.A., Naranjo, M.C.D., Daniilidou, M., Eiriksdottir, G., Mukherjee, S., Wallon, D., Uphill, J., Aspelund, T., Cantwell, L.B., Garzia, F., Galimberti, D., Hofer, E., Butkiewicz, M., Fin, B., Scarpini, E., Sarnowski, C., Bush, W.S., Meslage, S., Kornhuber, J., White, C.C., Song, Y., Barber, R.C., Engelborghs, S., Sordon, S., Voijnovic, D., Adams, P.M., Vandenberghe, R., Mayhaus, M., Cupples, L.A., Albert, M.S., De Deyn, P.P., Gu, W., Himali, J.J., Beekly, D., Squassina, A., Hartmann, A.M., Orellana, A., Blacker, D., Rodriguez-Rodriguez, E., Lovestone, S., Garcia, M.E., Doody, R.S., Munoz-Fernadez, C., Sussams, R., Lin, H., Fairchild, T.J., Benito, Y.A., Holmes, C., Karamujić-Čomić, H., Frosch, M.P., Thonberg, H., Maier, W., Roschupkin, G., Ghetti, B., Giedraitis, V., Kawalia, A., Li, S., Huebinger, R.M., Kilander, L., Moebus, S., Hernández, I., Kamboh, M.I., Brundin, R., Turton, J., Yang, Q., Katz, M.J., Concari, L., Lord, J., Beiser, A.S., Keene, C.D., Helisalmi, S., Kloszewska, I., Kukull, W.A., Koivisto, A.M., Lynch, A., Tarraga, L., Larson, E.B., Haapasalo, A., Lawlor, B., Mosley, T.H., Lipton, R.B., Solfrizzi, V., Gill, M., Longstreth, W.T., Montine, T.J., Frisardi, V., Diez-Fairen, M., Rivadeneira, F., Petersen, R.C., Deramecourt, V., Alvarez, I., Salani, F., Ciaramella, A., Boerwinkle, E., Reiman, E.M., Fievet, N., Rotter, J.I., Reisch, J.S., Hanon, O., Cupidi, C., Andre Uitterlinden, A.G., Royall, D.R., Dufouil, C., Maletta, R.G., de Rojas, I., Sano, M., Brice, A., Cecchetti, R., George-Hyslop, P.S., Ritchie, K., Tsolaki, M., Tsuang, D.W., Dubois, B., Craig, D., Wu, C.-K., Soininen, H., Avramidou, D., Albin, R.L., Fratiglioni, L., Germanou, A., Apostolova, L.G., Keller, L., Koutroumani, M., Arnold, S.E., Panza, F., Gkatzima, O., Asthana, S., Hannequin, D., Whitehead, P., Atwood, C.S., Caffarra, P., Hampel, H., Quintela, I., Carracedo, Á., Lannfelt, L., Rubinsztein, D.C., Barnes, L.L., Pasquier, F., Frölich, L., Barral, S., McGuinness, B., Beach, T.G., Johnston, J.A., Becker, J.T., Passmore, P., Bigio, E.H., Schott, J.M., Bird, T.D., Warren, J.D., Boeve, B.F., Lupton, M.K., Bowen, J.D., Proitsi, P., Boxer, A., Powell, J.F., Burke, J.R., Kauwe, J.S.K., Burns, J.M., Mancuso, M., Buxbaum, J.D., Bonuccelli, U., Cairns, N.J., McQuillin, A., Cao, C., Livingston, G., Carlson, C.S., Bass, N.J., Carlsson, C.M., Hardy, J., Carney, R.M., Bras, J., Carrasquillo, M.M., Guerreiro, R., Allen, M., Chui, H.C., Fisher, E., Masullo, C., Crocco, E.A., DeCarli, C., Bisceglio, G., Dick, M., Ma, L., Duara, R., Graff-Radford, N.R., Evans, D.A., Hodges, A., Faber, K.M., Scherer, M., Fallon, K.B., Riemenschneider, M., Fardo, D.W., Heun, R., Farlow, M.R., Kölsch, H., Ferris, S., Leber, M., Foroud, T.M., Heuser, I., Galasko, D.R., Giegling, I., Gearing, M., Hüll, M., Geschwind, D.H., Gilbert, J.R., Morris, J., Green, R.C., Mayo, K., Growdon, J.H., Feulner, T., Hamilton, R.L., Harrell, L.E., Drichel, D., Honig, L.S., Cushion, T.D., Huentelman, M.J., Hollingworth, P., Hulette, C.M., Hyman, B.T., Marshall, R., Jarvik, G.P., Meggy, A., Abner, E., Menzies, G.E., Jin, L.-W., Leonenko, G., Real, L.M., Jun, G.R., Baldwin, C.T., Grozeva, D., Karydas, A., Russo, G., Kaye, J.A., Kim, R., Jessen, F., Kowall, N.W., Vellas, B., Kramer, J.H., Vardy, E., LaFerla, F.M., Jöckel, K.-H., Lah, J.J., Dichgans, M., Leverenz, J.B., Mann, D., Levey, A.I., Pickering-Brown, S., Lieberman, A.P., Klopp, N., Lunetta, K.L., Wichmann, H.-E., Lyketsos, C.G., Morgan, K., Marson, D.C., Brown, K., Martiniuk, F., Medway, C., Mash, D.C., Nöthen, M.M., Masliah, E., Hooper, N.M., McCormick, W.C., Daniele, A., McCurry, S.M., Bayer, A., McDavid, A.N., Gallacher, J., McKee, A.C., van den Bussche, H., Mesulam, M., Brayne, C., Miller, B.L., Riedel-Heller, S., Miller, C.A., Miller, J.W., Al-Chalabi, A., Morris, J.C., Shaw, C.E., Myers, A.J., Wiltfang, J., O’Bryant, S., Olichney, J.M., Alvarez, V., Parisi, J.E., Singleton, A.B., Paulson, H.L., Collinge, J., Perry, W.R., Mead, S., Peskind, E., Cribbs, D.H., Rossor, M., Pierce, A., Ryan, N.S., Poon, W.W., Nacmias, B., Potter, H., Sorbi, S., Quinn, J.F., Sacchinelli, E., Raj, A., Spalletta, G., Raskind, M., Caltagirone, C., Bossù, P., Orfei, M.D., Reisberg, B., Clarke, R., Reitz, C., Smith, A.D., Ringman, J.M., Warden, D., Roberson, E.D., Wilcock, G., Rogaeva, E., Bruni, A.C., Rosen, H.J., Gallo, M., Rosenberg, R.N., Ben-Shlomo, Y., Sager, M.A., Mecocci, P., Saykin, A.J., Pastor, P., Cuccaro, M.L., Vance, J.M., Schneider, J.A., Schneider, L.S., Slifer, S., Seeley, W.W., Smith, A.G., Sonnen, J.A., Spina, S., Stern, R.A., Swerdlow, R.H., Tang, M., Tanzi, R.E., Trojanowski, J.Q., Troncoso, J.C., Van Deerlin, V.M., Van Eldik, L.J., Vinters, H. V., Vonsattel, J.P., Weintraub, S., Welsh-Bohmer, K.A., Wilhelmsen, K.C., Williamson, J., Wingo, T.S., Woltjer, R.L., Wright, C.B., Yu, C.-E., Yu, L., Saba, Y., Pilotto, A., Bullido, M.J., Peters, O., Crane, P.K., Bennett, D., Bosco, P., Coto, E., Boccardi, V., De Jager, P.L., Lleo, A., Warner, N., Lopez, O.L., Ingelsson, M., Deloukas, P., Cruchaga, C., Graff, C., Gwilliam, R., Fornage, M., Goate, A.M., Sanchez-Juan, P., Kehoe, P.G., Amin, N., Ertekin-Taner, N., Berr, C., Debette, S., Love, S., Launer, L.J., Younkin, S.G., Dartigues, J.-F., Corcoran, C., Ikram, M.A., Dickson, D.W., Nicolas, G., Campion, D., Tschanz, J., Schmidt, H., Hakonarson, H., Clarimon, J., Munger, R., Schmidt, R., Farrer, L.A., Van Broeckhoven, C., C. O’Donovan, M., DeStefano, A.L., Jones, L., Haines, J.L., Deleuze, J.-F., Owen, M.J., Gudnason, V., Mayeux, R., Escott-Price, V., Psaty, B.M., Ramirez, A., Wang, L.-S., Ruiz, A., van Duijn, C.M., Holmans, P.A., Seshadri, S., Williams, J., Amouyel, P., Schellenberg, G.D., Lambert, J.-C., Pericak-Vance, M.A. (2019) Genetic meta-analysis of diagnosed Alzheimer’s disease identifies new risk loci and implicates Aβ, tau, immunity and lipid processing. Nat. Genet. 51, 414–430.

Lee, B., Baek, J.-I., Min, H., Bae, S.-H., Moon, K., Kim, M.-A., Kim, Y.-R., Fogelgren, B., Lipschutz, J.H., Lee, K.-Y., Bok, J., Kim, U.-K. (2018) Exocyst Complex Member EXOC5 Is Required for Survival of Hair Cells and Spiral Ganglion Neurons and Maintenance of Hearing. Mol. Neurobiol. 55, 6518–6532.

Lee, M.-H., Lin, S.-R., Chang, J.-Y., Schultz, L., Heath, J., Hsu, L.-J., Kuo, Y.-M., Hong, Q., Chiang, M.-F., Gong, C.-X., Sze, C.-I., Chang, N.-S. (2010) TGF-β induces TIAF1 self-aggregation via type II receptor-independent signaling that leads to generation of amyloid β plaques in Alzheimer’s disease. Cell Death Dis. 1, e110.

McLaren, W., Gil, L., Hunt, S.E., Riat, H.S., Ritchie, G.R.S., Thormann, A., Flicek, P., Cunningham, F. (2016) The Ensembl Variant Effect Predictor. Genome Biol. 17, 122.

Mitchelmore, C., Büchmann-Møller, S., Rask, L., West, M.J., Troncoso, J.C., Jensen, N.A. (2004) NDRG2: a novel Alzheimer’s disease associated protein. Neurobiol. Dis. 16, 48–58.

Morris, L.G.T., Taylor, B.S., Bivona, T.G., Gong, Y., Eng, S., Brennan, C.W., Kaufman, A., Kastenhuber, E.R., Banuchi, V.E., Singh, B., Heguy, A., Viale, A., Mellinghoff, I.K., Huse, J., Ganly, I., Chan, T.A. (2011) Genomic dissection of the epidermal growth factor receptor (EGFR)/PI3K pathway reveals frequent deletion of the EGFR phosphatase PTPRS in head and neck cancers. Proc. Natl. Acad. Sci. U. S. A. 108, 19024–9.

Patel, D., Mez, J., Vardarajan, B.N., Staley, L., Chung, J., Zhang, X., Farrell, J.J., Rynkiewicz, M.J., Cannon-Albright, L.A., Teerlink, C.C., Stevens, J., Corcoran, C., Gonzalez Murcia, J.D., Lopez, O.L., Mayeux, R., Haines, J.L., Pericak-Vance, M.A., Schellenberg, G., Kauwe, J.S.K., Lunetta, K.L., Farrer, L.A. (2019) Association of Rare Coding Mutations With Alzheimer Disease and Other Dementias Among Adults of European Ancestry. JAMA Netw. Open 2, e191350.

Prekop, H.-T., Kroiss, A., Rook, V., Zagoraiou, L., Jessell, T.M., Fernandes, C., Delogu, A., Wingate, R.J.T. (2018) Sox14 Is Required for a Specific Subset of Cerebello–Olivary Projections. J. Neurosci. 38, 9539–9550.

Purcell, S., Neale, B., Todd-Brown, K., Thomas, L., Ferreira, M.A.R., Bender, D., Maller, J., Sklar, P., de Bakker, P.I.W., Daly, M.J., Sham, P.C. (2007) PLINK: a tool set for whole-genome association and population-based linkage analyses. Am. J. Hum. Genet. 81, 559–75.

Purcell, S.M., Wray, N.R., Stone, J.L., Visscher, P.M., O’Donovan, M.C., Sullivan, P.F., Sklar, P., Purcell Leader, S.M., Ruderfer, D.M., McQuillin, A., Morris, D.W., O’Dushlaine, C.T., Corvin, A., Holmans, P. a, Macgregor, S., Gurling, H., Blackwood, D.H.R., Craddock, N.J., Gill, M., Hultman, C.M., Kirov, G.K., Lichtenstein, P., Muir, W.J., Owen, M.J., Pato, C.N., Scolnick, E.M., St Clair, D., Sklar Leader, P., Williams, N.M., Georgieva, L., Nikolov, I., Norton, N., Williams, H., Toncheva, D., Milanova, V., Thelander, E.F., Sullivan, P.F., Kenny, E., Quinn, E.M., Choudhury, K., Datta, S., Pimm, J., Thirumalai, S., Puri, V., Krasucki, R., Lawrence, J., Quested, D., Bass, N., Crombie, C., Fraser, G., Leh Kuan, S., Walker, N., McGhee, K. a, Pickard, B., Malloy, P., Maclean, A.W., Van Beck, M., Pato, M.T., Medeiros, H., Middleton, F., Carvalho, C., Morley, C., Fanous, A., Conti, D., Knowles, J. a, Paz Ferreira, C., Macedo, A., Helena Azevedo, M., Kirby, A.N., Ferreira, M. a R., Daly, M.J., Chambert, K., Kuruvilla, F., Gabriel, S.B., Ardlie, K., Moran, J.L. (2009) Common polygenic variation contributes to risk of schizophrenia and bipolar disorder. Nature 10, 8192–8192.

Raghavan, N.S., Brickman, A.M., Andrews, H., Manly, J.J., Schupf, N., Lantigua, R., Wolock, C.J., Kamalakaran, S., Petrovski, S., Tosto, G., Vardarajan, B.N., Goldstein, D.B., Mayeux, R. (2018) Whole-exome sequencing in 20,197 persons for rare variants in Alzheimer’s disease. Ann. Clin. Transl. Neurol. 5, 832–842.

Rong, X.-F., Sun, Y.-N., Liu, D.-M., Yin, H.-J., Peng, Y., Xu, S.-F., Wang, L., Wang, X.-L. (2017) The pathological roles of NDRG2 in Alzheimer’s disease, a study using animal models and APPwt-overexpressed cells. CNS Neurosci. Ther. 23, 667–679.

Singhrao, S.K., Harding, A., Poole, S., Kesavalu, L., Crean, S. (2015) Porphyromonas gingivalis Periodontal Infection and Its Putative Links with Alzheimer’s Disease. Mediators Inflamm. 2015.

Song, W., Hooli, B., Mullin, K., Jin, S.C., Cella, M., Ulland, T.K., Wang, Y., Tanzi, R.E., Colonna, M. (2017) Alzheimer’s disease-associated TREM2 variants exhibit either decreased or increased ligand-dependent activation. Alzheimer’s Dement. 13, 381–387.

Subramanian, A., Tamayo, P., Mootha, V.K., Mukherjee, S., Ebert, B.L., Gillette, M.A., Paulovich, A., Pomeroy, S.L., Golub, T.R., Lander, E.S., Mesirov, J.P. (2005) Gene set enrichment analysis: a knowledge-based approach for interpreting genome-wide expression profiles. Proc Natl Acad Sci U S A 102, 15545–15550.

Tapia-Rojas, C., Inestrosa, N.C. (2018) Loss of canonical Wnt signaling is involved in the pathogenesis of Alzheimer’s disease. Neural Regen. Res. 13, 1705–1710. The PyMOL Molecular Graphics System, Version 2.4 (n.d.).

Tsavou, A., Curtis, D. (2019) In-silico investigation of coding variants potentially affecting the functioning of the glutamatergic N-methyl-D-aspartate receptor in schizophrenia. Psychiatr. Genet. 1.

Xiong, F., Ge, W., Ma, C. (2019) Quantitative proteomics reveals distinct composition of amyloid plaques in Alzheimer’s disease. Alzheimers. Dement. 15, 429–440.

Xu, J., Chatterjee, M., Baguley, T.D., Brouillette, J., Kurup, P., Ghosh, D., Kanyo, J., Zhang, Y., Seyb, K., Ononenyi, C., Foscue, E., Anderson, G.M., Gresack, J., Cuny, G.D., Glicksman, M.A., Greengard, P., Lam, T.T., Tautz, L., Nairn, A.C., Ellman, J.A., Lombroso, P.J. (2014) Inhibitor of the Tyrosine Phosphatase STEP Reverses Cognitive Deficits in a Mouse Model of Alzheimer’s Disease. PLoS Biol. 12, e1001923.

Zhang, X., Wu, Y., Cai, F., Song, W. (2019) Regulation of global gene expression in brain by TMP21. Mol. Brain 12, 39.

Zhang, Y., Kurup, P., Xu, J., Carty, N., Fernandez, S.M., Nygaard, H.B., Pittenger, C., Greengard, P., Strittmatter, S.M., Nairn, A.C., Lombroso, P.J. (2010) Genetic reduction of striatal-enriched tyrosine phosphatase (STEP) reverses cognitive and cellular deficits in an Alzheimer’s disease mouse model. Proc. Natl. Acad. Sci. 107, 19014–19019.

Zhou, R., Yang, G., Guo, X., Zhou, Q., Lei, J., Shi, Y. (2019) Recognition of the amyloid precursor protein by human γ-secretase. Science (80-.). 363, eaaw0930.

Zhou, X., Edmonson, M.N., Wilkinson, M.R., Patel, A., Wu, G., Liu, Y., Li, Y., Zhang, Z., Rusch, M.C., Parker, M., Becksfort, J., Downing, J.R., Zhang, J. (2016) Exploring genomic alteration in pediatric cancer using ProteinPaint. Nat. Genet. 48, 4–6.

